# Cognitive Vergence and Pupillary Responses as Functional Oculomotor Signatures to Differentiate AT(N) Biological Profiles

**DOI:** 10.64898/2026.04.14.718456

**Authors:** Ricardo Martínez-Flores, Isabel Martín-Sobrino, Neus Falgàs, Oriol Grau-Rivera, Marc Suárez-Calvet, Carlos Cristi-Montero, Agustín Ibañez, Hans Supèr

## Abstract

**Background:** The AT(N) biological framework classifies Alzheimer’s disease (AD) pathology using CSF biomarkers, with the A+T+ profile defining biological AD and the A−T+ profile representing a biologically distinct entity consistent with suspected non-Alzheimer’s pathophysiology, such as primary age-related tauopathy. Functional assessment capable of differentiating these profiles non-invasively remains limited. This study investigates whether cognitive vergence and pupillary temporal dynamics during a visual oddball task can distinguish A−T+ from A+T+ biological profiles in individuals with mild cognitive impairment (MCI).

**Methods:** Thirty-eight participants with MCI (12 A−T+, 26 A+T+) classified by CSF biomarkers completed a visual oddball task (80% distractors, 20% targets) under continuous eye-tracking. Linear mixed-effects models examined profile × condition interactions on full time series and six trial-level temporal features. Participant-level differentiation was assessed using binomial logistic regression, adjusting for age, sex, and MMSE.

**Results:** Both profiles showed comparable overall oculomotor response magnitudes but diverged markedly in temporal organization. Significant profile × condition interactions emerged for cognitive vergence global slope, time to peak, and pupillary time to peak. Logistic regression confirmed that timing features discriminated biological profiles at the participant level, with differentiation reversing direction between distractor and target conditions. A−T+ participants also maintained superior target detection accuracy (89.3% vs. 82.4%, p = 0.001).

**Conclusion:** Cognitive Vergence and pupillary temporal dynamics during an oddball task provide condition-dependent functional oculomotor signatures that systematically differentiate AT(N) biological profiles in MCI, suggesting that oculomotor assessment may offer an accessible, non-invasive complement to CSF-based profile characterization.

## 1. Introduction

Alzheimer’s disease (AD) is a neurodegenerative condition driven by the progressive accumulation of specific proteinopathies that can be detected and biologically staged during life through validated biomarkers [1]. The societal impact of AD is substantial: an estimated 50 million people are currently affected worldwide, a number projected to surpass 115 million by 2050 as populations age [2]. Crucially, the neuropathological cascade underlying AD unfolds years before cognitive symptoms become apparent [1], which has historically constrained opportunities for early research and therapeutic intervention. In response to these challenges, the 2018 NIA-AA Research Framework established a biological rather than syndromal definition of AD, organizing biomarkers into the AT(N) classification system according to pathological process: amyloid-β (A), fibrillar tau (T), and neurodegeneration (N) [1]. The recently revised 2024 Alzheimer’s Association criteria further refined this framework by differentiating Core 1 biomarkers (early-changing phosphorylated tau fragments (p-tau181, p-tau217, p-tau231) and amyloid markers) from Core 2 biomarkers reflecting tau aggregate burden, and by anchoring AD diagnosis to abnormality on Core 1 biomarkers [3]. Across both frameworks, the A+T+ profile biologically defines AD, characterized by reduced CSF Aβ42 and elevated phosphorylated tau, while the A−T+ profile—tau-positive in the absence of detectable amyloid pathology—represents a biologically distinct entity whose etiology may primarily reflect primary age-related tauopathy (PART) or other non-Alzheimer tauopathies [4,5].

Both A+T+ and A−T+ profiles share the presence of tau-related pathology, and post-mortem evidence consistently identifies the locus coeruleus (LC) as the earliest site of tau accumulation in the AD pathological cascade, preceding cortical involvement by years [4,6].

Located in the pontine brainstem, the LC constitutes the principal noradrenergic nucleus, with extensive projections to cortical and subcortical structures implicated in arousal, attentional regulation, and oculomotor control [7–9]. Through direct anatomical connections with the Edinger-Westphal nucleus, LC activity influences pupillary responses [7,10]. Regarding vergence, the LC sends noradrenergic projections to the superior colliculus [11], a structure in which distinct neuronal populations encode and modulate vergence eye movements [12], suggesting an indirect pathway by which LC activity may influence vergence control. Pupil diameter serves as a well-validated psychophysiological index of LC-noradrenergic function across multiple cognitive paradigms [7,10], while cognitive vergence responses correlate with attentional allocation and visual event-related potentials in parietal regions [13,14]. Given the shared embryological origin of ocular and neural tissues [15], cognitive vergence and pupillary assessment during structured cognitive tasks may offer a non-invasive window onto the functional integrity of neural systems affected early by tau pathology in both biological profiles.

Prior work has documented that cognitive vergence and pupillary alterations in individuals with MCI and AD during oddball paradigms, demonstrating that both signals are sensitive to attentional network dysfunction associated with cognitive decline [16]. Building on this, previous work demonstrated that CSF biomarkers, Aβ42, p-Tau181 and Aβ42/p-Tau ratio, significantly modulate oculomotor responses during oddball tasks, with these responses predicting tau burden independently of age, sex, and MMSE [17]. However, a critical question remains unaddressed: whether oculomotor signatures differ specifically between biological profiles, that is, whether the presence of amyloid co-pathology in A+T+ individuals generate a qualitatively distinct oculomotor pattern compared to A−T+ individuals. This distinction is biologically meaningful because A+T+ and A−T+ profiles differ not only in amyloid status but in the degree to which cortical network coordination may be compromised. In A+T+, amyloid accumulation preferentially affects default mode network (DMN) regions including the posterior cingulate cortex and precuneus [18,19], potentially disrupting the anti-correlated relationship between task-positive and task-negative networks necessary for efficient attentional engagement [20]. A−T+ individuals, spared from this amyloid-driven network disruption, may retain relative functional resilience under high attentional demand. Consistent with this view, previous literature reported that A−T+ older adults exhibited significantly greater hippocampal volume, cortical thickness across AD-associated regions, and cerebral glucose metabolism compared to A+T+ individuals, while remaining indistinguishable from cognitively unimpaired healthy A-T-controls on all neurodegenerative markers [21], suggesting that tau protein in the absence of amyloid does not produce the same degree of structural and metabolic compromise. Whether these differences in pathological profile translate into detectable differences in task-evoked oculomotor dynamics has not been examined.

This question carries direct clinical relevance. Profile characterization under both the 2018 NIA-AA framework and the revised 2024 criteria relies on CSF extraction or PET imaging, procedures that are invasive, costly, and poorly suited to population-level monitoring or routine longitudinal follow-up [18]. Although blood-based biomarkers such as plasma p-tau217 offer a less invasive alternative with increasing clinical availability, they share with CSF and PET the limitation of providing static indices of pathological burden without capturing how the brain responds functionally to cognitive demand. Thus, oculomotor assessment during structured attentional tasks could complement existing biomarker approaches by providing an accessible, non-invasive measure of functional network integrity, particularly informative in contexts where pathological burden and functional capacity may dissociate.

Building on these considerations, the present cross-sectional study aims to determine whether cognitive vergence and pupillary response patterns in individuals with MCI classified according to AT(N) biological profiles can functionally differentiate the A−T+ profile from the A+T+ profile. By establishing these relationships within the AT(N) framework, we seek to advance the validation of cognitive vergence and pupillary responses as accessible functional oculomotor signatures that complement CSF-based assessments by providing insight into the patient’s cerebral functionality, particularly the attentional and oculomotor networks, thus bridging biological and functional manifestations for early identification and monitoring of different neuropathological profiles.

## 2. Methods

A cross-sectional design was used in this study. Ethical approval was obtained from the Ethics Committees of the University of Barcelona (HCB/2021/0668) and was conducted in accordance with the Declaration of Helsinki. The study followed the STROBE reporting guidelines for cross-sectional studies [22], and the corresponding checklist is provided as supplementary material.

### 2.1 Participants

Thirty-eight individuals in MCI stage (26 women, 12 men) from two Barcelona-based hospitals (Hospital del Mar and Hospital Clínic) participated in this study. All participants underwent lumbar puncture for CSF extraction as part of their clinical evaluation. Based on these clinical assessments, all participants showed abnormal CSF core AD biomarkers in accordance with Alzheimer’s Association criteria [3].

Written informed consent was obtained from all participants or their legal representatives prior to enrollment. Participants and caregivers received comprehensive verbal and written instructions regarding all experimental procedures.

### 2.2 Biological Profile Classification

Biological profiles were established following the AT(N) framework [1] using population-specific CSF biomarker thresholds validated for Spanish cohorts. Hospital del Mar participants were classified using CORCOBIA study thresholds [23]: Aβ42 <750 pg/mL defined amyloid positivity (A+), pTau >69.85 pg/mL indicated tau positivity (T+), and tTau >522.0 pg/mL marked neurodegeneration (N+). For Hospital Clínic participants, institutional validated cutoffs were applied: Aβ42 ≤600 pg/mL (A+), pTau >65 pg/mL (T+), and tTau >385 pg/mL (N+). Despite the use of center-specific thresholds, the same classification procedure and analytical approach were applied uniformly across participants. Hospital center was included as a covariate in all statistical models; (see statistical analysis section), it did not reach statistical significance in any analysis, suggesting that center-related variability did not systematically influence the results.

Based on these cut-off points, participants were stratified into two primary biological profiles: the A+T+ profile, characterized by concurrent amyloid and phosphorylated tau abnormalities, and the A-T+ profile, defined by tau abnormalities in the absence of amyloid pathology.

Given the exploratory nature of this study and limited sample size (n=38), our classification approach incorporated additional considerations. Two participants presenting A−T−(N)+ profiles (isolated elevation of total tau) were grouped with the A-T+ profile, justified by established evidence demonstrating total tau as a sensitive marker across various tauopathies and an important biomarker of neurofibrillary degeneration [5,24].

To further assess the robustness of this decision, a sensitivity analysis was conducted excluding these two participants (see statistical analysis and supplementary material) and did not meaningfully alter the pattern of findings. This methodological approach aligns with the AT(N) framework’s explicit recommendation for adaptation “when it is fit for the purpose of specific research goals” [1,3], maintaining consistency with NIA-AA principles while enabling meaningful biological stratification.

CSF concentrations of Aβ42, tTau, and pTau were quantified using single molecule array technology (Lumipulse platform) following standardized institutional protocols [25] at each participant’s respective hospital.

### 2.3 Sample size and statistical power

Given that participant enrollment was contingent on the availability of CSF biomarker data (requiring lumbar puncture as part of a separate clinical protocol) rather than guided by formal power calculations, we performed a post-hoc simulation-based power analysis to evaluate the adequacy of our sample for detecting the effects of interest.

Standard power formulas are not appropriate for hierarchical time-series data, as they assume independence between observations. To account for the nested structure of our dataset (repeated temporal measurements within participants) and the random effects specification of our models (participant-level intercepts and time slopes), we used a parametric bootstrap approach, which is more appropriate than analytical formulas for mixed-effects models with this type of data structure [26]. Specifically, we generated 1,000 new datasets by drawing from the estimated parameter distributions of the fitted mixed-effects model and refitting the full model to each simulated dataset with SIMR r package approach [27].

Applied to the strongest interaction observed in our analyses (cognitive vergence response as a function of biological profile × stimulus condition; β = 0.096, SE = 0.011, t = 8.99, p < .001), this procedure yielded a post-hoc power estimate of 100% (95% CI: [99.6%, 100%]) at α = .05, with all 1,000 simulated datasets producing p < .001. The high detection rate reflects the combination of a large effect size and the precision afforded by the dense temporal sampling inherent to eye-tracking paradigms.

We further evaluated how power varied as a function of sample size by repeatedly subsampling participants (n = 20, 25, 30, 35) and rerunning the simulation procedure. Power was essentially at ceiling for samples of 25 or more participants (> 99%). At n = 20, estimated power averaged 90%, though with considerable variability across draws (range: 56%–100%), indicating that with very small samples the ability to detect this interaction becomes sensitive to the specific composition of the group. Taken together, these results support the adequacy of the current sample for the primary analyses reported.

### 2.4 Inclusion/exclusion Criteria

Participants were eligible for enrollment if they met the following criteria: (1) age ≥65 years; (2) availability of recent CSF biomarker analysis (Aβ42, p-Tau, t-Tau) obtained through lumbar puncture as part of clinical evaluation; (3) adequate visual acuity (with or without corrective lenses) to perceive on-screen stimuli; and (4) capacity to provide informed consent or availability of a legal representative to provide consent on behalf of the participant.

Participants were excluded based on the following criteria: (1) severe cognitive impairment (Mini-Mental State Examination score <10); (2) documented neurological conditions with clinically significant cognitive impact (e.g., cerebrovascular disease); (3) severe psychiatric diagnoses; (4) structural brain abnormalities identified through neuroimaging that could contribute to cognitive dysfunction (e.g., intracranial neoplasms); (5) ophthalmological conditions interfering with task performance (including blindness, strabismus, nystagmus, or significant retinal/oculomotor pathology); and (6) insufficient Spanish language comprehension.

### 2.5 Eye-Tracking Equipment

Ocular data acquisition utilized the BGaze system (BraingazeSL, Mataró, Spain), which integrates stimulus presentation with binocular eye position and pupillometry recording. Visual stimuli were displayed at 1024 × 768-pixel resolution. Eye movements were captured using a Tobii 5L remote eye tracker (Tobii Technology AB, Sweden) sampling at 33 Hz. Manufacturer specifications report spatial accuracy of 0.4° visual angle and precision of 0.32° visual angle under optimal binocular viewing conditions. Random measurement noise inherent to eye-tracking data is effectively minimized through trial averaging [28].

### 2.6 Procedure

Testing occurred in a dedicated room within the hospital facility under controlled lighting conditions (lights extinguished, curtains closed to achieve dim ambient illumination 50 lux on average). Participants were positioned approximately 50 cm from the stimulus display screen, with the eye-tracker mounted directly below. Corrective lenses were permitted. Head position was stabilized using a chinrest to minimize motion artifact.

### 2.7 Task Design

We employed a visual oddball paradigm, selected for its robust activation of distributed brain networks (https://neurosynth.org/analyses/terms/oddball/) including the locus coeruleus [29]. The task comprised 120 discrete trials, each beginning with a 2000 ms gray mask screen, followed by 2000 ms presentation of a centrally positioned letter string (Figure 1).

**Figure 1.**
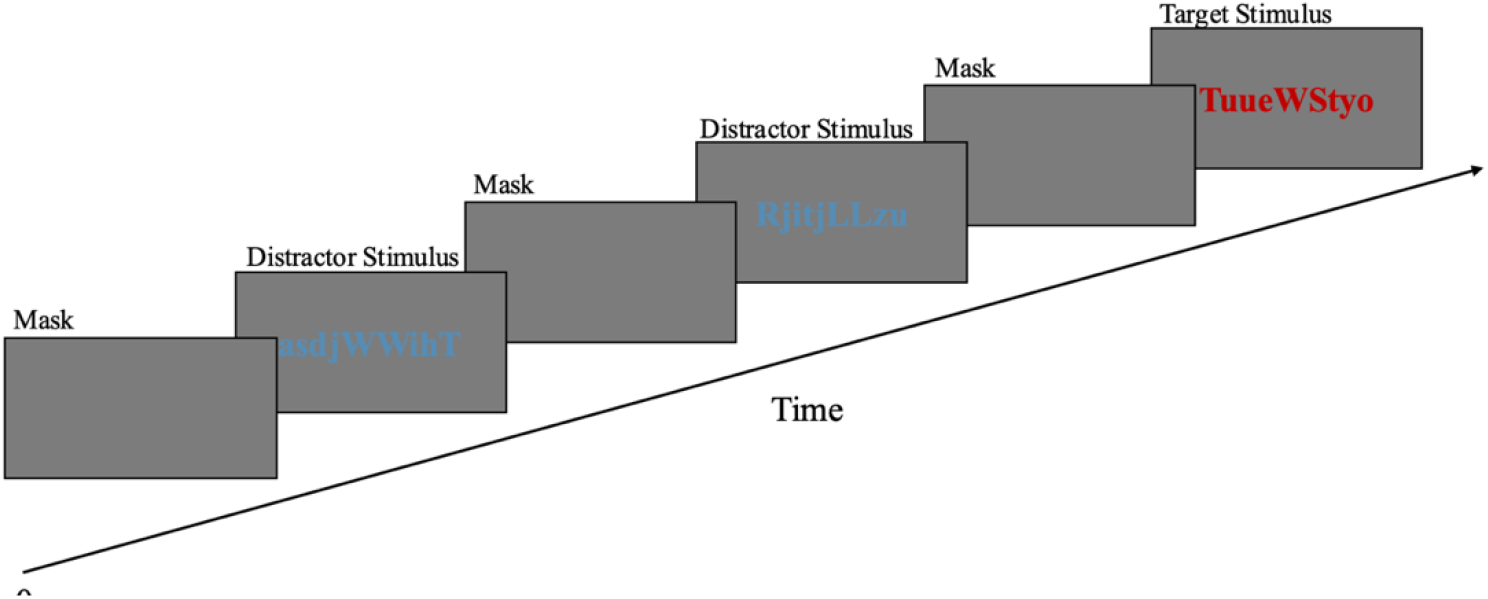
Oddball Task secuence. Strings of symbols are presented for two seconds, followed by a grey mask. In 80% of the trials the symbols were printed in blue, and in the remaining 20% they were printed in red.

Each stimulus consisted of 11 randomly generated characters in mixed upper and lower case, arranged to avoid forming meaningful words or recognizable acronyms that might introduce cognitive bias. Stimuli were differentiated solely by font color: 80% appeared in blue (frequent/non-target condition) while 20% appeared in red (rare/target condition). Participants received instructions to maintain visual fixation on the screen and execute a button press exclusively upon detecting red-colored strings. Verbal instruction reinforcement was provided by the experimenter as needed during task performance.

Following standard oddball nomenclature, red letter strings constituted target stimuli while blue strings served as distractor stimuli. Stimulus presentation order was pseudorandomized. Continuous binocular eye-tracking was maintained throughout the ∼6-minute task duration. Prior to task initiation, a five-point binocular calibration procedure was completed for each participant to ensure measurement accuracy.

### 2.8 Signal Preprocessing and Functional Oculomotor Signal Extraction

Cognitive vergence and pupil responses were extracted during the attentional task as oculomotor signature. Eye-tracking recordings underwent systematic preprocessing to ensure signal integrity. Quality control procedures excluded trials containing: (1) fixations outside the display area (normalized coordinates <0 or >1); (2) physiologically implausible vergence angles (<0° or >20°); and (3) rapid eye movements exceeding 150 mm/s velocity threshold. Trials were retained only if valid data comprised at least 30% of the recording period, ensuring adequate signal-to-noise ratios for subsequent analyses. After excluding trials with more than 30% missing samples, the remaining trials showed limited need for reconstruction: 71.8% required ≤20% interpolation and 28.2% required <5%, consistent with expected blink-related signal interruptions in a clinical population, consistent with established eye-tracking methodologies [30].

Vergence angle was computed using a vector-based geometric approach from three-dimensional eye position coordinates. Two-dimensional gaze coordinates were first converted to physical dimensions (millimeters) using the monitor’s specifications (340 mm × 190 mm). Gaze direction vectors were constructed from each eye’s spatial position to its corresponding point of regard on the screen. Vergence angle (θ) at each time point was calculated as the angle between left and right gaze vectors using vector dot product geometry, expressed in degrees thus relative vergence changes were computed as υ(t) = (γ(t) - γ□), where γ□ represents baseline vergence, this method was previously published by our team [14]. For pupil responses, absolute diameter changes were calculated as π(t) = p(t) - p□, where p(t) represents the mean of left and right pupil diameters. Both baseline values (γ□ and p□) were calculated as the mean of the first 7 temporal points (≈212 ms pre-stimulus period) following previous recommendations [30]. Both cognitive vergence and pupil signals were subjected to Gaussian smoothing using moving average filter prior to baseline correction to optimize signal-to-noise ratio.

To quantify the functional characteristics of ocular responses beyond simple amplitude measures, six features were extracted from each individual trial’s vergence and pupil response curves. Initial slope was computed over the first 500 ms, capturing the primary response dynamics. Global slope characterized the overall rate of change across the entire trial duration. Late slope was computed over 1000-2000 ms, reflecting sustained response or plateau phase. Temporal windows for slope extraction were determined through visual inspection of mean response curves. Linear regression was employed for slope estimation as it provides a robust single-parameter summary of the dominant directional trend within each temporal window, minimizing the influence of high-frequency noise while capturing the primary rate of signal change. Area under the curve (AUC) was computed using trapezoidal integration, the standard approach in pupillometry [30], representing the cumulative magnitude of the response. Time to peak identified the latency to maximum response amplitude. Peak amplitude measured the maximum value attained during the trial. Slope values were converted from milliseconds to seconds (multiplied by 1000) for interpretability. These features were extracted separately for both cognitive vergence and pupil responses, yielding 12 total features per trial (6 for each measure). All features were subsequently standardized (z-scores) prior to statistical modeling to enable direct comparison across measures with different units and scales. All the preprocessing and features extractions was conducted in Python 3.x.

### 2.8 Statical Analysis

Data normality was assessed using Shapiro-Wilk tests and visual inspection with histograms and Q-Q plots. While pupil data exhibited normal distribution, cognitive vergence data appeared approximately normal in histograms but showed slight tail deviations in Q-Q plots. Given the repeated-measures nature of eye-tracking data and the large dataset (237,600 data points; 6,600 per participant), linear mixed-effects models were employed, as they are robust to minor deviations from normality in large samples [31]. Because different CSF cut-off values were applied across recruitment centers, center was initially evaluated as a fixed covariate, however, it showed no significant effects in any model specification. Following the principle of parsimony and in the absence of strong theoretical justification relative to other covariates, the center variable was not retained in final models.

Our analytical approach adopted a hierarchical strategy to characterize biological profile differences at multiple levels of temporal and spatial aggregation. Because the data consist of high-resolution temporal sequences nested within trials and participants, traditional multiple comparison corrections such as Bonferroni or FDR are not appropriate, as they assume independent observations. Instead, we adopted the principle of “keep it maximal” to prevent type I error by including the necessary random effects to appropriately model the hierarchical structure and temporal dependencies [32]. This hierarchical approach included: (a) models using the full cognitive vergence and pupil time series as dependent variables with Timestamp as a random slope to capture participant-specific temporal dynamics, (b) trial-level analyses examining whether biological profiles differed in task performance accuracy, (c) decomposition into trial-level ocular biomarkers (temporal slope, timing, and magnitude biomarkers) to explore whether biological profiles differed in temporal dynamics versus response magnitude, and (d) participant-level analyses testing whether ocular biomarkers showing significant profile differences could distinguish between biological profiles using binomial logistic regression.

All models were adjusted for age, sex, and MMSE. Across analyses, Profile distinguished A−T+ from A+T+ biological profiles (with A+T+ as the reference category), Condition was coded with distractor as the reference category, and the Profile × Condition interaction tested whether stimulus differentiation varied between biological profiles.

For the full time-series analyses, cognitive vergence and pupil responses (standardized as Z-scores) were analyzed using linear mixed-effects models including random intercepts for Participant and participant-specific random slopes for timestamp to account for individual temporal trajectories.

Task performance accuracy was analyzed at the trial level using linear mixed-effects models with random intercepts for Participant; because these models did not retain within-trial temporal resolution, timestamp was not included as a random slope.

To dissociate temporal dynamics from response magnitude, oculomotor signals were decomposed into trial-level ocular biomarkers as described in the methods section 2.8. Each standardized ocular biomarker was analyzed separately using linear mixed-effects models with random intercepts for Participant. As these analyses operated on trial-level summaries, the within-trial temporal structure was no longer present and thus did not require a random slope for timestamp.

Oculomotor signature showing significant Profile × Condition interactions were subsequently aggregated at the participant level (averaged within condition and re-standardized) and analyzed using separate binomial logistic regression models predicting biological profile membership. At this aggregation level, each participant contributed one observation per condition, eliminating the hierarchical trial structure and therefore not requiring random effects. Odds ratios (OR) and 95% confidence intervals were derived from the fitted models.

To assess the robustness of our findings, a sensitivity analysis was conducted by excluding two A-T+ participants who were classified based on total tau rather than ptau181 values. All three analytical levels (full time series, trial-level features, and participant-level classification) were re-estimated with this restricted sample. Comparison between primary and sensitivity analyses revealed that cognitive vergence time to peak showed a marginal reduction in significance (p = 0.067), while all other significant effects exhibited slight variations in magnitude but maintained statistical significance below the α = 0.05 (see supplementary material, tables S3 and S4). Significance was set at p < 0.05 for all analyses. All analyses were conducted in R (version 4.x).

## 3. Results

The two biological profiles were well-matched on demographic and cognitive characteristics, with no significant differences in age, sex distribution, or general cognitive performance (Table 1). As expected by design, the groups diverged markedly on Aβ42 and Aβ42/p-Tau ratio, confirming successful stratification of participants into biologically distinct profiles.

**Table 1.** Participants Characterization.

| Variable | All<br>n = 38 | A-T+<br>n = 12 | A+T+<br>n = 26 | p value<br>(A-T+ vs A+T+) |
| --- | --- | --- | --- | --- |
| Age (y) | 72.9 ± 5.83 | 73.4 ± 4.93 | 72.7 ± 6.28 | 0.975 |
| Sex (W/M) | 26/12 | 9/3 | 17/9 | 0.559 |
| MMSE | 22.4 ± 4.04 | 23.8 ± 3.93 | 21.7 ± 4.00 | 0.154 |
| Aβ42 | 623 ± 281 | 946 ± 255 | 473 ± 123 | **<0.001** |
| p-Tau | 160 ± 196 | 97.6 ± 70.7 | 131 ± 84.7 | 0.120 |
| t-Tau | 762 ± 370 | 754 ± 273 | 824 ± 373 | 0.172 |
| Aβ42/p-Tau ratio | 7.80 ± 7.77 | 15.1 ± 10.6 | 4.6 ± 2.21 | **<0.001** |

Data is presents for continuous variables as mean ± standard deviation and for categorical variables (sex) as frequency. (y): years; Sex (W/M): Women/Male; P-value from t-test (chi-square for sex). In bold significant p value.

### 3.1 Functional Oculomotor Signatures: cognitive vergence and pupil response

Both cognitive vergence and pupillary signals showed systematic modulation by stimulus type across the full sample, with target stimuli consistently eliciting larger responses than distractors (Figure 2). Neither cognitive vergence nor pupillary amplitude differed between profiles when averaged across conditions. However, significant interactions between stimulus type and biological profile emerged for both signals (Table 2), indicating that the magnitude of condition-dependent modulation varied between the two groups.

**Table 2.** Global and temporal analysis of cognitive vergence and pupil.

| Signal | Effect | $\beta$ | SE | p |
| --- | --- | --- | --- | --- |
| **Cognitive Vergence** | Condition | 0.039 | 0.0056 | **<0.001** |
|  | Profile | -0.074 | 0.0841 | 0.386 |
|  | Interaction | 0.096 | 0.0107 | **<0.001** |
| **Pupil** | Condition | 0.467 | 0.0054 | **<0.001** |
|  | Profile | 0.007 | 0.0620 | 0.916 |
|  | Interaction | -0.060 | 0.0103 | **<0.001** |

Reference categories: Condition = Distractor; Profile = A+T+. Bold values indicate $p < 0.05$ . $\beta$ = standardized regression coefficient; SE = standard error.

**Figure 2.**
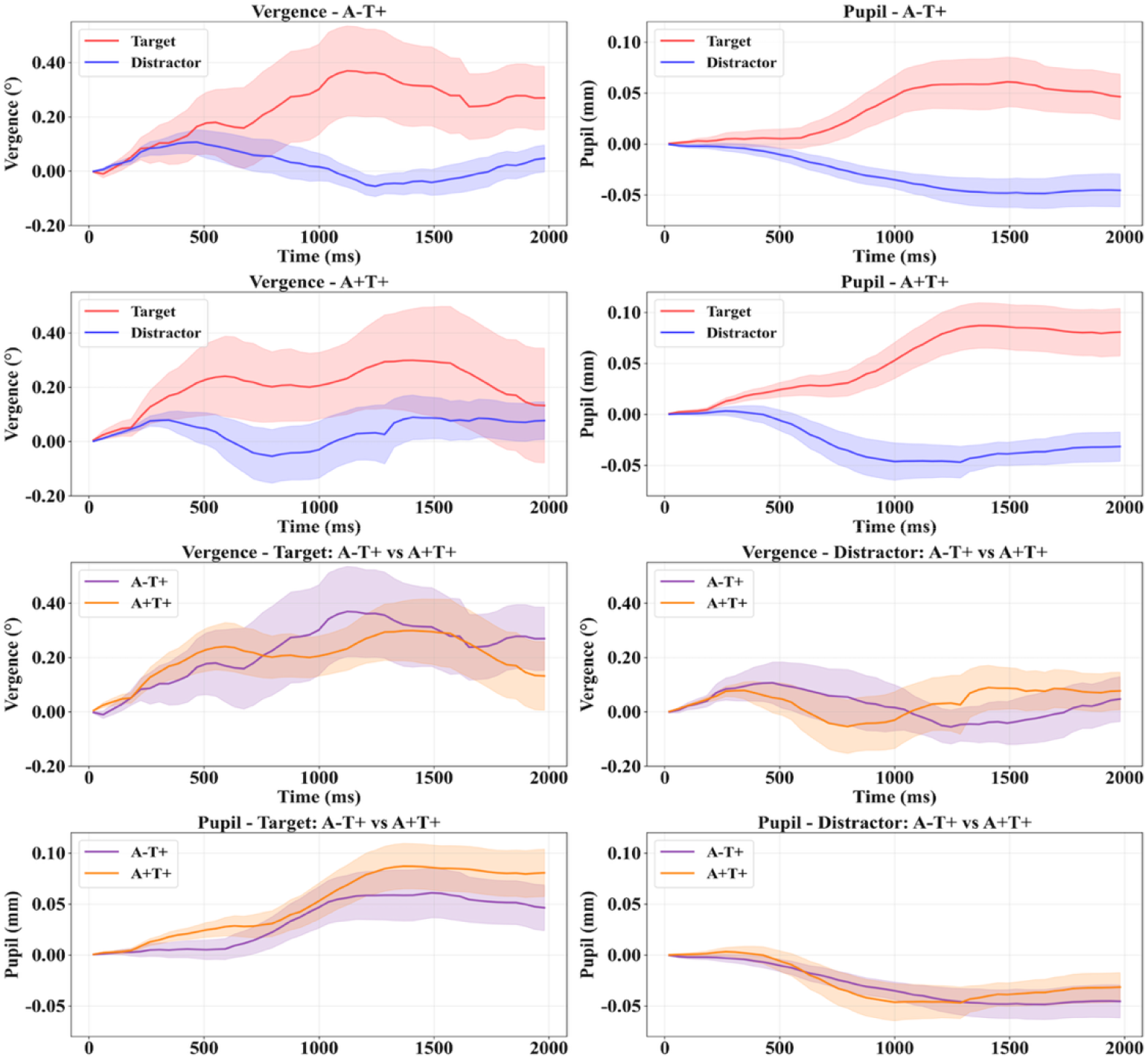
Cognitive vergence and pupil response during oddball task. Rows 1-2: Average cognitive vergence and pupil responses during oddball target/distractor conditions according to biological profile. Rows 3-4: Target vs. distractor comparisons by biological profile. Shaded areas = standard error; time = milliseconds from stimulus onset.

The pattern of this modulation differed between the two ocular systems. For cognitive vergence responses, the A-T+ profile showed greater target-distractor differentiation compared to the A+T+ profile. For pupillary responses, the A+T+ profile exhibited greater pupillary dilation compared to the A-T+ profile. These divergent interaction patterns demonstrate that the two biological profiles generated distinct oculomotor response signatures depending on stimulus type, with each profile showing differential engagement of cognitive vergence versus pupillary systems.

### 3.2 Accuracy Response

Behavioral performance revealed profile-dependent differences that varied by stimulus condition (Figure 3). Both groups performed at comparable levels on distractor trials, with near-ceiling accuracy. However, performance patterns diverged on target trials, where the A-T+ profile maintained higher accuracy while the A+T+ profile showed reduced performance.

**Figure 3.**
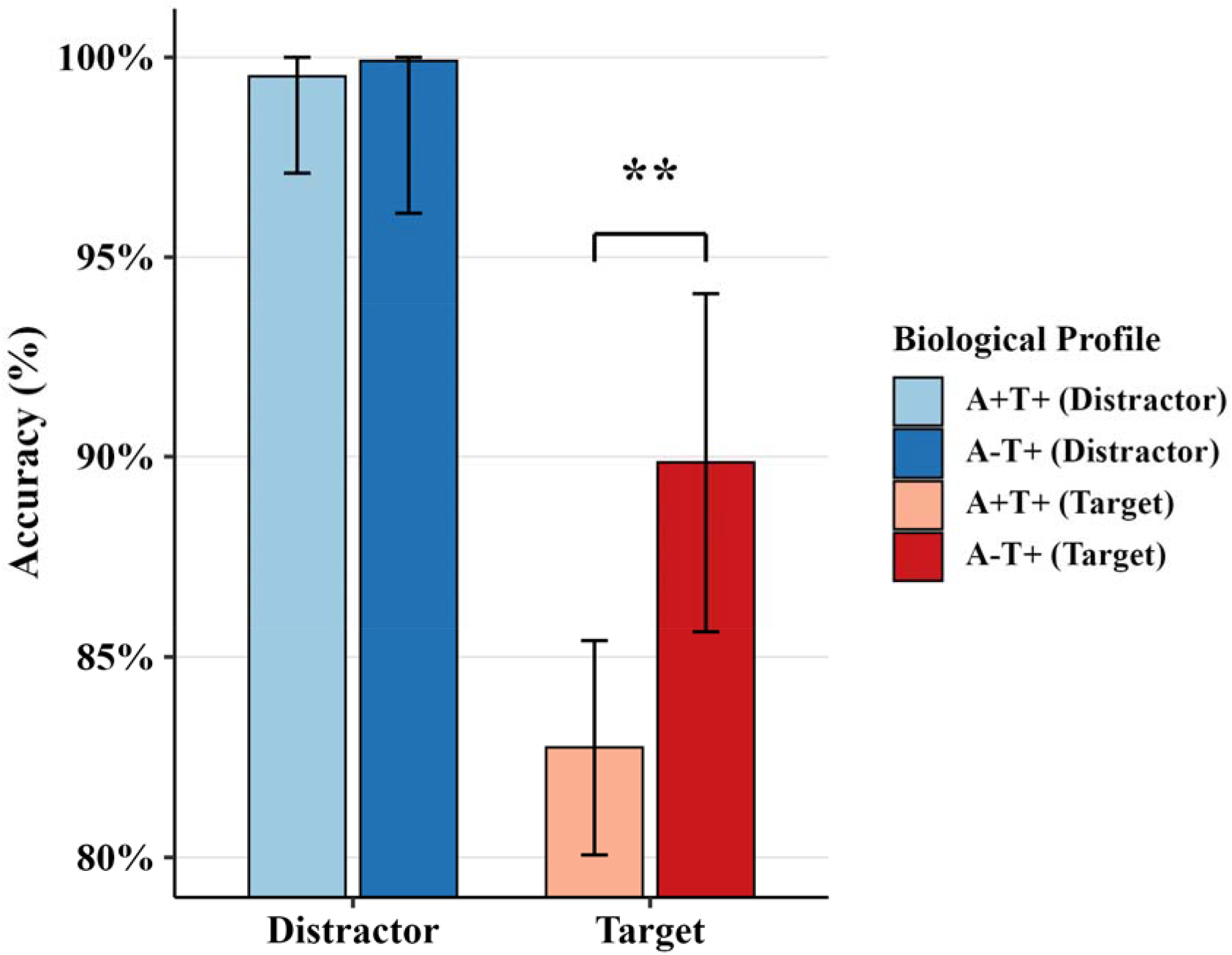
Accuracy comparison to oddball task. Accuracy in percentage, asterisk indicates significant difference between profiles.

### 3.3 Functional Oculomotor Signatures: trial-level features

Trial-level feature analysis revealed condition-dependent oculomotor signatures that systematically differentiated A-T+ from A+T+ biological profiles (Table 3). Linear mixed-effects models identified significant profile-by-condition interactions for vergence global slope, vergence time to peak, and pupillary time to peak. For vergence global slope, A-T+ exhibited steeper vergence dynamics than A+T+ specifically during Target trials, with no difference between profiles during Distractor trials. For vergence time to peak, A-T+ showed later vergence peaks than A+T+ during Distractor trials, a pattern that reversed during Target trials, where A+T+ exhibited the slower response. Pupillary time to peak followed the same directional pattern and additionally showed a significant profile main effect, indicating that A-T+ had globally later pupillary peaks than A+T+ across conditions, with the between-profile difference being largest during Distractor trials and attenuating during Target trials.

**Table 3.**
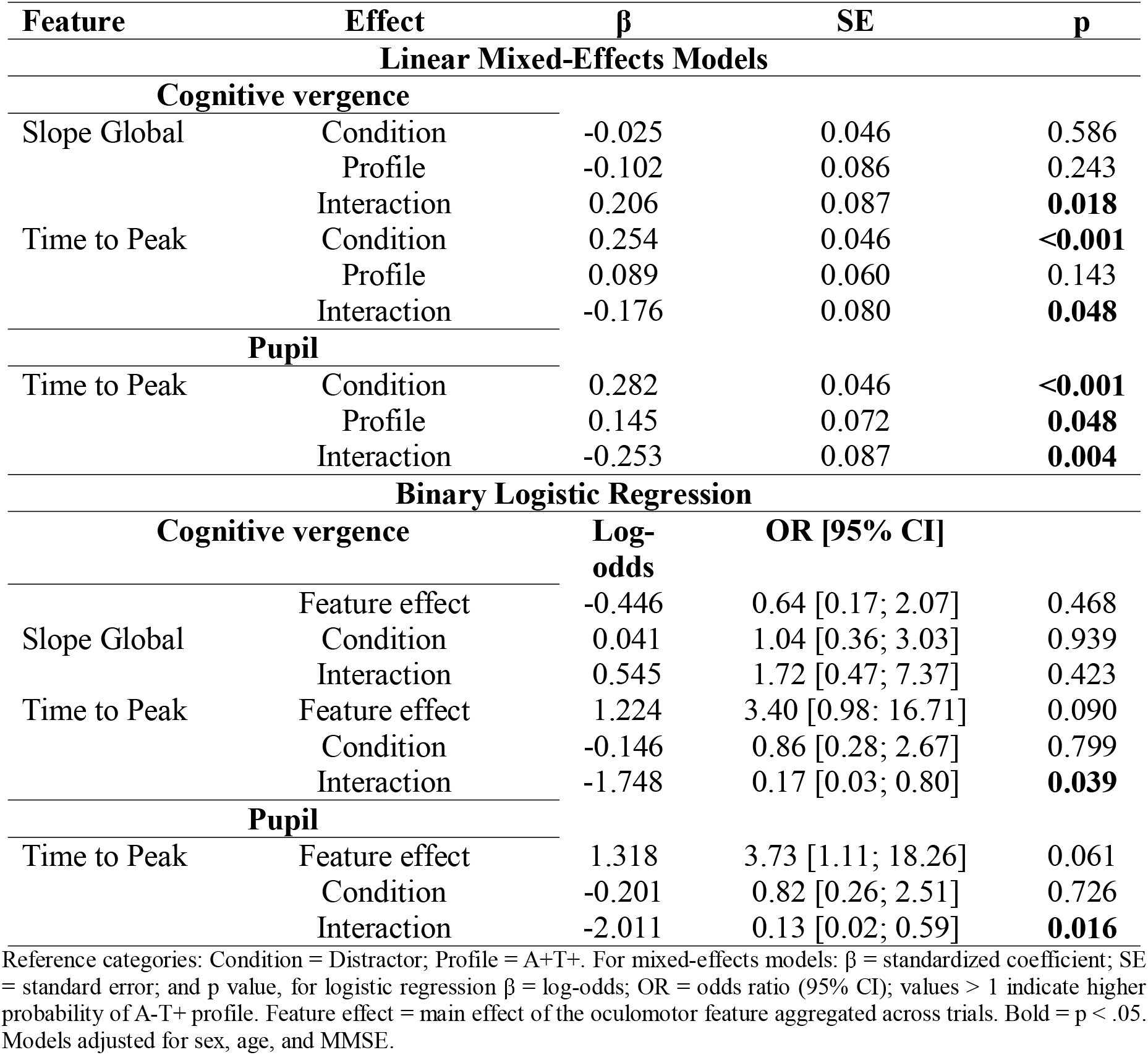
Condition-Dependent Temporal Oculomotor Features Differentiating AT(N) Biological Profiles.

Binary logistic regression confirmed that this condition-dependency extended to the classification of biological profile at the subject level. For vergence time to peak, the discriminative capacity of the feature was significantly greater during Distractor than Target trials, with later vergence peaks predicting A-T+ profile membership in the Distractor condition. Pupillary time to peak showed the same pattern with greater effect magnitude: later pupillary peaks were associated with higher odds of A-T+ profile during Distractor trials, and this relationship was significantly attenuated during Target trials.

## 4. Discussion

This study aimed to determine whether cognitive vergence and pupillary response patterns in individuals with MCI classified according to AT(N) biological profiles can functionally differentiate the A−T+ profile from the A+T+ profile. The findings consistently indicate that the two profiles are not distinguished by the overall magnitude of their oculomotor responses, but rather by the temporal organization of those responses, specifically by when peak responses occurs and that this temporal differentiation is condition-dependent, varying systematically between distractor and target trials. This pattern was observed across analytical levels, from continuous signal dynamics to trial-level feature decomposition and subject-level analyses, lending convergent support to the main observation.

### 4.1 Condition-dependent temporal dissociation between biological profiles

The condition-dependent nature of the observed dissociation is central to interpreting these findings. Both profiles showed comparable oculomotor response magnitudes overall, yet their timing signatures diverged in a direction that depended on whether the stimulus was a distractor or a target. For both cognitive vergence and pupillary time to peak, A-T+ showed later responses under distractor conditions, while A+T+ showed the most delayed timing under target conditions. This reversal implies that the two profiles differ not in the degree of oculomotor slowing but in the conditions under which slowing is expressed, suggesting that qualitatively distinct neural processes may underlie the timing differences in each profile. The behavioral findings parallel this pattern: both groups performed comparably on distractor trials, where accuracy was near ceiling, while A-T+ maintained higher target detection accuracy than A+T+ specifically on target trials. This convergence between behavioral and oculomotor condition-dependency strengthens the interpretation that the observed differences are functionally meaningful rather than incidental variability.

### 4.2 Neural mechanisms underlying profile-specific oculomotor timing

These results may possibly be explained by the differential impact that isolated tau versus combined amyloid-tau pathology has on the neural systems that govern the temporal dynamics of attentional processing. Both profiles carry tau pathology, and post-mortem evidence consistently identifies the LC as the earliest site of tau accumulation in the AD pathological cascade, preceding cortical involvement by years [4,6]. Through its widespread noradrenergic projections, the LC modulates arousal, attentional state, and oculomotor output via direct connections to the superior colliculus and the Edinger-Westphal nucleus [7,9]. Critically, the LC operates in two functionally distinct modes: tonic firing that regulates baseline attentional state and phasic bursting that is selectively evoked by salient stimuli such as oddball targets [33]. Pupil diameter has been established as a reliable psychophysiological index of LC activity in both its tonic and phasic modes, with trial-level associations between pupillary responses and LC neural activity demonstrated in oddball paradigms specifically [10,29,33]. Under distractor conditions, which impose minimal attentional demand and engage primarily tonic noradrenergic regulation, the globally delayed pupillary time-to-peak observed in the A-T+ group, including a significant profile main effect, may reflect subtle alterations in tonic LC-mediated modulation associated with tau burden at this structure. Tau pathology in the LC is known to disrupt both tonic and phasic noradrenergic output; early LC tau burden shifts neuronal activity toward a state of persistent high tonic discharge that secondarily impairs the efficiency of phasic bursting [34]. Under this tonic-phasic imbalance, such differences in tonic regulation would be most detectable under low-demand conditions, where LC tonic output constitutes the primary driver of attentional state and compensatory network recruitment is not available to mask the underlying dysfunction [29,35],

Under target conditions, successful oculomotor modulation requires more than tonic LC regulation. Oddball target processing robustly engages distributed attentional networks, including the dorsal attention network implementing top-down selection, the ventral attention network mediating bottom-up reorientation to salient stimuli, the frontoparietal network coordinating executive control and decision-related processing and the hippocampus, which supports attention-dependent memory encoding [29,29,36–38]. The phasic LC response to target stimuli has been specifically linked to the P300 event-related potential and to the encoding of salience information through gamma power increases in prefrontal cortex [33,36]. For this distributed network response to unfold efficiently, appropriate deactivation of the DMN is also required: regions that overlap substantially with DMN nodes are among the earliest sites of amyloid accumulation, including the posterior cingulate cortex and precuneus [19,39], and higher amyloid burden has been associated with aberrant DMN connectivity even in cognitively unimpaired individuals [19,20,39]. Failure of appropriate DMN suppression during goal-directed processing may disrupt the anti-correlated relationship between task-positive and task-negative networks that is necessary for efficient attentional engagement [20,29,40]. In A+T+, where amyloid pathology is superimposed on tau burden, this additional disruption of cortical network coordination may explain why target processing, which places the greatest demand on this coordinated network response, produces the most delayed oculomotor timing in this profile. The integrity of frontoparietal connectivity has been specifically linked to attentional resilience in the context of early LC tau pathology, with higher left frontoparietal network connectivity attenuating the negative effects of LC tau burden on cognitive performance [41]. In A-T+, where amyloid disruption is absent, this network-level resilience may be better preserved, which would be consistent with the observed maintenance of target detection accuracy and the absence of abnormally delayed target-related timing in this profile. This interpretation is structurally supported by evidence that A-T+ older adults show significantly greater hippocampal volume and cortical thickness across AD-associated regions compared to A+T+ individuals, while remaining neurodegeneration ally indistinguishable from cognitively unimpaired healthy controls [21], suggesting that the relative functional advantage observed here in oculomotor timing may reflect a genuinely preserved neural substrate rather than mere compensatory recruitment.

### 4.3 Selectivity and robustness of oculomotor timing features

Among all features extracted, only timing metrics (time to peak for both cognitive vergence and pupil) consistently differentiated the profiles across both trial-level and subject-level analyses. Slope-based metrics capturing the rate of signal change showed more limited and less stable discrimination. This selectivity is consistent with prior evidence that peak latencies of both cognitive vergence and pupil responses are sensitive to attentional network dysfunction in cognitive impairment [16], and further extends this observation by showing that even within a cognitively impaired sample, timing features carry information about the specific molecular pathology combination present. Time to peak integrates information across the full trial window, reflecting the cumulative temporal unfolding of the response rather than summarizing a single phase, which may account for its greater robustness [42]. The selective failure of slope metrics at the subject level, despite their significance at the trial level for cognitive vergence global slope, suggests that condition-dependent slope differences are real but insufficiently stable within individuals to serve as reliable subject-level markers, whereas timing metrics appear to capture more stable individual-level characteristics of temporal processing.

The differential robustness of cognitive vergence and pupillary time to peak also warrants comment. Pupillary time to peak showed significant effects at all levels of analysis, including a profile main effect reflecting globally later peaks in A-T+ independent of condition. Cognitive vergence time to peak showed only the condition-dependent interaction without a profile main effect. This difference in sensitivity is broadly consistent with the anatomical organization of these systems: pupillary modulation is more directly coupled to LC activity through both sympathetic and parasympathetic pathways and has been validated as a psychophysiological index of LC-noradrenergic function across multiple paradigms [7,10,35], while vergence responses depend additionally on circuits involved in spatial attention and binocular processing that introduce condition-specific influences [9,14,16]. These systems are not independent, however: vergence movements are known to trigger pupillary responses through direct anatomical coupling at the Edinger-Westphal nucleus, which receives input from vergence-related circuits and coordinates both oculomotor outputs [7–9]. Under this coupling, cognitive vergence responses temporally precede and may condition the subsequent pupillary response, such that the greater robustness of pupillary timing across conditions could partly reflect the integration of vergence-driven input with direct LC-noradrenergic modulation at the Edinger-Westphal level [7,9]. The fact that LC neurons phase-lock to prefrontal and hippocampal oscillatory rhythms that synchronize to behavioral events [8,37,38] further supports the idea that pupillary timing captures a more direct and condition-independent readout of noradrenergic state, whereas vergence timing may reflect additional demand-dependent processes that emerge under attentional conditions.

The sensitivity analysis excluding two participants classified on the basis of total tau rather than ptau181 confirmed robustness, with all major findings maintained and only minor variations in effect magnitude. This supports interpretability within the AT(N) framework as applied, while acknowledging that measurement heterogeneity within the A-T+ group warrants attention in future work with larger samples.

### 4.4 Clinical implications

A distinctive feature of the present findings is that cognitive vergence and pupillary temporal dynamics appear to reflect not merely which molecular pathology profile is present, but the functional state of the neural systems that profile compromises. The condition-dependent pattern of oculomotor timing differences captures how each profile processes attentional demands differently, providing a dynamic readout of brain network integrity rather than a static proxy for pathological load. This distinction is clinically relevant: current AT(N) profile characterization relies on CSF extraction or PET imaging, procedures that are invasive, costly, and difficult to implement at scale for population monitoring or longitudinal follow-up [18]. Although blood-based biomarkers such as ptau-217 represent a less invasive alternative that is increasingly available in clinical settings, they share with CSF and PET the limitation of providing a static index of pathological burden without capturing how the brain responds functionally to cognitive demand. Oculomotor assessment during structured attentional tasks could complement these approaches precisely in this regard, offering an accessible, non-invasive measure of functional brain integrity that may be particularly informative in contexts where pathological burden and functional capacity dissociate [15,43]

The observation that A-T+ profiles maintained better target detection accuracy and relatively preserved oculomotor timing under high-demand conditions, despite carrying tau pathology, is consistent with evidence that molecular pathology alone does not determine functional outcomes and that network-level compensatory capacity plays an important role during early disease stages [41,43]. This raises the possibility that temporal oculomotor metrics could serve as accessible indicators of functional reserve and may be particularly relevant for monitoring treatment response in interventions targeting noradrenergic function or attentional network synchronization, where changes in functional capacity may precede detectable changes in static biomarkers [44]. The differential oculomotor signatures between profiles also suggest that these populations may respond differently to network-targeted interventions, a question that warrants direct investigation in future trials.

### 4.5 Strength and limitations

This study represents, to our knowledge, the first systematic description of cognitive vergence and pupillary temporal features as functional oculomotor signatures differentiating biological profiles defined by AT(N) criteria during an attentional task. The hierarchical analytical strategy, spanning continuous signal dynamics, trial-level feature decomposition, and subject-level analyses, provides a multi-resolution characterization that is more informative than any single level alone. The large number of temporal observations per participant supports the reliability of the extracted features, and the consistency of findings across analytical approaches strengthens confidence in the main observations. The use of AT(N) biological profile classification based on CSF markers, rather than clinical diagnosis, aligns with current frameworks and reduces the confounding influence of cognitive heterogeneity on oculomotor findings.

Important limitations must be acknowledged. The sample size is small, constraining statistical power and limiting generalizability. The cross-sectional design precludes any inference about whether oculomotor timing differences precede, accompany, or follow specific stages of pathological accumulation. The absence of a cognitively unimpaired control group prevents characterization of normative oculomotor timing in this paradigm, which would be needed to determine whether the observed patterns represent deviations from typical aging or reflect pathology-specific alterations. Longitudinal designs with larger and more homogeneously characterized samples, including cognitively unimpaired individuals at preclinical stages, are needed to determine whether these temporal oculomotor signatures track pathological progression and whether they are detectable prior to cognitive impairment.

## Conclusion

This study demonstrates that cognitive vergence and pupillary temporal dynamics during an oddball task provide condition-dependent oculomotor signatures that systematically differentiate A-T+ from A+T+ biological profiles. The discriminative information resides in the timing of peak responses rather than their magnitude, and the direction of profile differences reverses depending on whether stimuli require passive processing or active attentional engagement. These results may possibly reflect the differential impact of isolated tau pathology versus combined amyloid-tau pathology on LC-mediated tonic modulation and cortical attentional network recruitment, with each profile showing its functional vulnerability under the conditions that most tax the neural systems it primarily compromises. Ocular biomarkers represent a promising and accessible complement to CSF-based profile characterization, and their validation in longitudinal and preclinical samples constitutes a meaningful next step.

## Acknowledgment

R.M-F was supported by a grant from the National Agency for Research and Development (ANID)/Scholarship Program/DOCTORADO BECAS CHILE/2024–(Grant Nº 72240103. NF was the recipient of the Juan Rodés contract JR22/00014 (Instituto de Salud Carlos III, Spain), Alzheimer’s Association (AACSF_21_723056 to NF) and Instituto de Salud Carlos III Spain, and co-funded by the European Union (PI25/00222 to NF). AI is supported by grants from the Multi-partner consortium to expand dementia research in Latin America [ReDLat, supported by Fogarty International Center (FIC), National Institutes of Health, National Institutes of Aging (R01 AG057234, R01 AG075775, R01 AG21051, R01 AG083799, CARDS-NIH, R01 AG057234), Alzheimer’s Association (SG-20-725707), Rainwater Charitable Foundation – The Bluefield project to cure FTD, and Global Brain Health Institute)], ANID/FONDECYT Regular (1250091 and 1210176 and 1220995); ANID/PIA/ANILLOS ACT210096; JPI JPND-Care, DISCeRN 2025 - Health and Social Care Research with a Focus on the Moderate and Late Stages of Neurodegenerative Diseases; FONDEF ID20I10152, and ANID/FONDAP 15150012; Wellcome Trust award for BRAIN-CLIMA: Investigating the Combined Impact of Heat and Air Pollution on Blood-Brain Barrier Integrity and Brain Aging in Latin America, (335293/Z/25/Z), and the CliCBrain (Horizon ID: 101236426; DOI 10.3030/101236426, Marie Skłodowska-Curie Actions - MSCA). The contents of this publication are solely the responsibility of the authors and do not represent the official views of these institutions. The funders had no role in study design, data collection and analysis, decision to publish or preparation of the manuscript.

## Data availability statement

Data is available via request to the corresponding author. The method of cognitive vergence for measuring neurodegenerative proteins has been protected by a patent application.

## Declaration of generative AI and AI-assisted technologies in the writing process

During the preparation of this work the author(s) used ChatGPT-4o in order to improve language and readability. After using this tool/service, the author(s) reviewed and edited the content as needed and take(s) full responsibility for the content of the publication.

## Patient consent statement

All participants provided informed written consent to participate in the study.

